# Non-Homology-Based Prediction of Gene Functions

**DOI:** 10.1101/730473

**Authors:** Xiuru Dai, Zheng Xu, Zhikai Liang, Xiaoyu Tu, Silin Zhong, James C. Schnable, Pinghua Li

## Abstract

Advances in genome sequencing and annotation have eased the difficulty of identifying new gene sequences. Predicting the functions of these newly identified genes remains challenging. Genes descended from a common ancestral sequence are likely to have common functions. As a result homology is widely used for gene function prediction. This means functional annotation errors also propagate from one species to another. Several approaches based on machine learning classification algorithms were evaluated for their ability to accurately predict gene function from non-homology gene features. Among the eight supervised classification algorithms evaluated, random forest-based prediction consistently provided the most accurate gene function prediction. Non-homology-based functional annotation provides complementary strengths to homology-based annotation, with higher average performance in Biological Process GO terms, the domain where homology-based functional annotation performs the worst, and weaker performance in Molecular Function GO terms, the domain where the accuracy of homology-based functional annotation is highest. Non-homology-based functional annotation based on machine learning may ultimately prove useful both as a method to assign predicted functions to orphan genes which lack functionally characterized homologs, and to identify and correct functional annotation errors which were propagated through homology-based functional annotations.

## Introduction

The rapid acceleration in genome sequencing is providing complete sequences for dozens of new species each year (Michael and Jackson, 2013; Chen et al., 2018). Advances in both *de novo* and extrinsic evidence based gene structure annotation, combined with low cost and abundant RNA sequence datasets, aid the identification and definition gene models across each new genome assembly (Campbell et al., 2014; Del Angel et al., 2018; Cook et al., 2019; Monnahan et al., 2019). However, while the accuracy and throughput of methods to define the structure of genes have grown rapidly, methods to experimentally determine the function of individual genes have not. Existing annotations are taken from a small set of proteins with direct experimental evidence and then these annotations are extrapolated to not only homologous genes in the same genome but homologous genes in the genomes of other species (Valencia, 2005). Among eukaryotes, fission yeast *Schizosaccharomyces pombe* has perhaps the most comprehensive set of functional gene annotations (Aslett and Wood, 2006). There are currently 41,912 gene associations for 5,397 gene products available on *Sz. pombe* GeneDB (Lock et al., 2018). Of these, 16,657 functional annotations for 2,302 genes (42.6% of 5,397 annotated genes) are directly derived from experiments. Of those, a subset of 4,761 functional annotations for 1,459 genes (27.0% of all annotated gene models in *Sz. pombe*) are supported by mutant phenotype analysis. Among flowering plants, the model species *Arabidopsis thaliana* has been the subject of intensive and comprehensive genetic investigation. However, of the 28,775 annotated gene models in the TAIR10 *A. thaliana* reference genome, only 14.2% have functional annotations supported by mutant phenotypes, 25.8% have functional annotations supported by other types of experimental evidence (e.g. biochemical assays of enzymatic function, protein-protein interactions, expression data, etc), 51.1% are functionally annotated based on solely protein features, sequence similarity, or other homology-based forms of evidence, and 8.9% of gene models lack any functional annotation (Lamesch et al., 2011).

Two decades ago, a significant challenge of homology-based functional annotation, as generally employed, was recognized: functional annotations are propagated from one sequence to the next without data on provenance. Thus it is often impossible or impractical to track a computationally assigned functional annotation back to the original source of experimental evidence. This presents a challenge, as mistaken findings related to protein functions will be published from time to time (Iyer et al., 2001), and once an experimentally derived functional annotation is assigned to homologs in other species, there is no way to “recall” this annotation. In fact, the annotation is likely to continue to propagate to new genome assemblies and to reannotations of existing assemblies (Brenner, 1999; Valencia, 2005; Gilks et al., 2002, 2005). Curated annotations made based on non-sequence similarity evidence have been estimated to have an error rate of 13%-18%, while curated annotations made based on sequence similarity evidence had an estimated error rate of 49% (Jones et al., 2007). In short, “functional annotations are propagated repeatedly from one sequence to the next, to the next, with no record made of the source of a given annotation, leading to a potential transitive catastrophe of erroneous annotations” (Karp, 1998). Homology-based functional annotation also rests on the basic assumption that sequence similarity and functional similarity is highly correlated, which is an assumption that is not always correct as demonstrated by many cases of sub- and neo- functionalization between homologous genes (Clark and Radivojac, 2011; Brown et al., 2006; Radivojac et al., 2013).

In addition to concerns with annotation accuracy, many species also contain a signficant number of genes where homology-based annotation is not possible. The genomes of arabidopsis and rice, respectively, are reported to contain 1,430 and 1,926 orphan genes which lack known homologs in other species (Guo et al., 2007; Guo, 2013). By definition, homology-based methods are only able to make predictions when the function of at least one related sequence – whether detected through direct nucleotide or protein sequence similarity (Conesa et al., 2005), or more sensitive methods such as the presence of a shared protein domain or protein domain architecture (Thomas et al., 2003; Quevillon et al., 2005; Hulo et al., 2006; Finn et al., 2015) – has been experimentally characterized. As the result, genes belonging to orphan gene families and/or carrying only domains of unknown functions are likely to lack predicted or potential functions. This in turn contributes to the noted pattern of clustering of research efforts on more detailed characterization of genes with existing well characterized functions (Stoeger et al., 2018).

However, there exists a parallel set of nonhomology based approaches to predict the function of uncharacterized genes (Marcotte et al., 1999; Marcotte, 2000; Gabaldón and Huynen, 2004). Chromosomal context has been widely employed for functional prediction in prokaryotes where operons of genes involved in a single metabolic pathway or biological process are common (Edwards et al., 2005; Enault et al., 2005). High rates of gene loss and horizontal gene transfer in prokaryotes can also be employed to assign predicted functions to genes with either similar or complementary phylogenetic distributions (Gaasterland and Ragan, 1998; Pellegrini et al., 1999; Morett et al., 2003). In eurkaryotes such as maize and arabidopsis, mRNA co-expression analysis has been show to improve the prioritization of GWAS hits (Chan et al., 2011; Angelovici et al., 2017; Schaefer et al., 2018; Zheng et al., 2019). Protein co-expression networks are also beginning to become more widely available and appear to capture different information content from mRNA co-expression networks (Walley et al., 2016). Nonhomology based methods have been used to systematically develop functional predictions in prokaryotes and have been to prioritize individual sets of candidate genes in plants (Chan et al., 2011; Angelovici et al., 2017; Schaefer et al., 2018; Zheng et al., 2019). However, genome-wide functional annotation in plants still relies primarily on homology-based methods.

This initial analysis of the potential for non-homology-based genome-wide functional annotation in eukaryotes focuses on maize (*Zea mays*), a widely studied genetic model and economically vital crop species. As the result, maize has an extensive collection of functional genomic datasets, including large RNA and protein expression atlases (Walley et al., 2016; Stelpflug et al., 2016), methylation and histone modification profiling datasets (Dong et al., 2017), one of the largest collection of characterized and cloned loss of function mutants of any plant species (Schnable and Freeling, 2011; Oellrich et al., 2015). However, maize also exhibits substantial genomic structural complexity with thousands of genes which are present in the genomes and genome assemblies of some inbreds but not others (Springer et al., 2009; Swanson-Wagner et al., 2010; Sun et al., 2018; Springer et al., 2018). The majority of these genes are evolutionarily young, with the evidence that the majority of them are conserved between maize and its wild progenitor teosinte, but only a small minority were found to be conserved at syntenic locations the genome of *Sorghum bicolor* a related species which diverged from the lineage leading to maize approximately 12 million years ago (Swigonňová et al., 2004; Sun et al., 2018; Springer et al., 2018). The need for non-homology-based functional annotations is pressing in maize, particularly as there is evidence these new and variably-present genes may be involved in hybrid vigor (Paschold et al., 2014; Baldauf et al., 2016, 2018). In this project, we evaluated the potential for using supervised machine learning-based classification algorithms to predict the function of annotated maize genes using purely non-homolog-based predictive variables.

## Results

We assembled a set of descriptors for each gene, including gene structure, population genetics, expression, histone modification and DNA methylation features (Table S1). This dataset included features calculated from the alignment of sequence reads to the maize AGPv4 reference genome and data mined from previously published papers (Jiao et al., 2017; Hoopes et al., 2019; Walley et al., 2016; Dong et al., 2017; Bukowski et al., 2017; Liang et al., 2019). Some important features, for example, protein abundance data for maize vegetative and reproductive stages, are only available for prior versions of the maize reference genome. As the result, we constrained this analysis and only considered a set of 29,428 gene models which had a 1:1 relationship between a single gene model in the maize reference genome version AGPv2 and a single gene model in the maize reference genome version AGPv4 (Schnable et al., 2009; Wang et al., 2016; Liang et al., 2019). Many algorithms for making predictions from input feature sets are intolerant of missing values. While the overall rate of missing data for this 1:1 gene set was low, a small number of genes (1,995 genes) have missing values for more than half of the total set of 369 features (Figure S1). These genes were omitted from subsequent analyses. For the remaining 27,433 genes, missing values were imputed using the median value for that feature across all the genes where that feature was successfully scored.

### Potential for Dimensional Reduction Among Nonhomology Features

Spearman correlation coefficients were caclulated among the 369 features (Figure 1). The two largest classes of features, i.e. RNA abundance (274 features) and protein abundance (33 features), showed substantial between-feature correlation. Supervised machine learning classification models tend to overfit when trained with excessively large numbers of features. This overfitting decreases predictive performance on non-training datasets. Dimensional reduction techniques seek to to address this problem by reducing the total number of features available for training without significantly reducing the overall information content of the dataset. Dimensional reduction algorithms can generally be divided into the categories of feature extraction and feature selection. Principal Component Analysis (PCA) is a widely used feature extraction technique that improves learning performance, reduces computational complexity, builds better models and decreases the required memory space (Tang et al., 2014). More than 90% of the variance in RNA abundance and protein abundance could be captured by 20 and 10 principal components respectively (Figure 1). These principal components were used in place of the original RNA and protein abundance features, which decreased the number of possible predictive variables from 369 to 92. This decrease also substantially reduced the degree of correlations between possible predictive variables. The 95^th^ percentile and 99^th^ percentile for the absolute values of Spearman correlation (r_s_) dropped from 0.76 and 0.94 to 0.22 and 0.51, respectively (Figure 1B).

**Figure 1:**
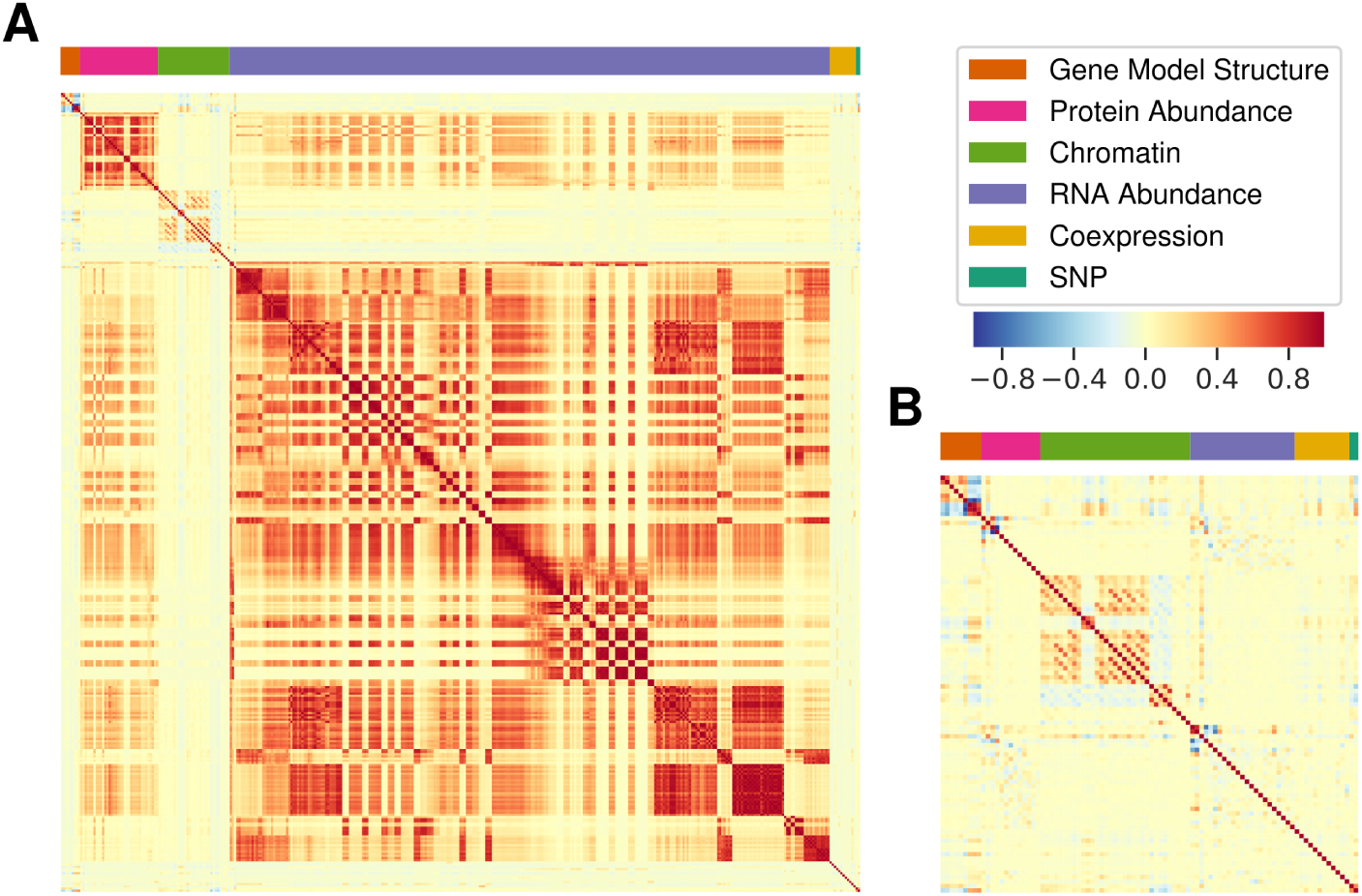
Correlations among different features in the prediction dataset. (A) Spearman correlations and group membership among all 369 original features. (B) Spearman correlations and group membership for 92 features remaining after targeted principal component-based dimension reduction of mRNA and protein-based expression data. The ordering of individual features from top to bottom and left to right is provided in Table S2.

Multiple distinct sets of GO predictions exist for the maize reference genome (Goodstein et al., 2011; Tello-Ruiz et al., 2015; Wimalanathan et al., 2018). We chose to use the maize GAMER dataset as our starting point for training and evaluating non-homology-based prediction algorithms (Wimalanathan et al., 2018). The published maize GAMER dataset includes 9,336 GO terms which are directly assigned to one or more gene models, and an additional 2,757 GO terms which are implicitly assigned to one or more gene models. An implicit GO term assignment can occur when a specific GO term is explicitly assigned to a gene. In this case, each parent of that GO term is also assigned to the same gene implicitly. We utilized both implicit and explicit GO term assignments. The initial dataset thus consisted of 12,093 GO terms and each go term was assigned to one or more of the 39,324 gene models in the B73 AGPv4 maize reference genome. We chose to exclude both extremely common GO terms (e.g. GO:0008150 “Biological Process”) and extremely rare GO terms. Extremely common GO terms tend to be low information content. Extremely rare GO terms are unlikely to possess enough known positive genes to accurately train prediction algorithms. After excluding GO terms assigned to *<*100 genes in our set of 27,433 genes with feature data and those assigned to >5,000 genes in the same set, 1,562 GO terms – including 1,148 Biological Process, 151 Cellular Component and 263 Molecular Function terms – remained for downstream analyses.

Classifiers were trained using either all 369 predictor features or the reduced set of 92 predictor features remaining after targeted dimensional reduction. Across the models trained of each of the 1,562 GO terms in this analysis, those trained with the full set of 369 features exhibited prediction accuracies of 0.35 to 0.93 with a median of 0.67. Models trained using the reduced set of 92 predictor features exhibited prediction accuracies of 0.41 to 0.93 with a median of 0.68. The increase in prediction accuracy for the targeted dimensional reduction models relative to the full models was statistically significant (p=0.0002; two tailed paired t-test). While this targeted dimensional reduction increased prediction accuracy, untargeted dimensional reduction did not. Models trailed using a set of 50 principal components extracted from the complete set of 369 features provided significantly lower prediction accuracy than either the total feature set (p = 5.96e-158; two tailed paired t-test) or the targeted dimensional reduction feature set (p = 3.89e-192; two tailed paired t-test) (Figure S2). Prediction accuracy was evaluated using shuffled data to test whether, even after dimension reduction, over fitting might be occurring. Prediction accuracy balanced gene sets with shuffled functional annotations ranged from 0.47 to 0.57 with a median of 0.51, only slightly higher than expected of random predictions. All subsequent analyses used the set of 92 features obtained by targeted dimension reduction of protein and RNA abundance features.

### Selection of Random Forest for Gene Function Prediction

Eight machine learning-based supervised classification algorithms including random forest (Liaw and Wiener, 2002), stochastic gradient boosting machines (Ridgeway et al., 2013), Lasso and Elastic-Net Regularized Generalized Linear Models (Friedman et al., 2010; Simon et al., 2011), radial kernel SVM (support vector machine) (Karatzoglou et al., 2004, 2018), partial least squares (Wehrens and Mevik, 2007), neural network (Venables and Ripley, 2002; Ripley et al., 2016), penalized multinomial regression (Venables and Ripley, 2002; Ripley et al., 2016), and linear discriminant analysis (Venables and Ripley, 2002; Ripley et al., 2013) were evaluated for their accuracy in predicting GO annotations. Benchmark genes for every GO terms were divided into the sets of 80% training and 20% testing. For training data, 10-fold cross validation was performed for all machine learning methods. Validation accuracy and testing accuracy, i.e. the accuracy in testing dataset, were both calculated for the 8 algorithms and comparisons of the algorithms were based on the accuracy in testing dataset. Based on the average accuracy across all GO terms tested, random forest and gbm methods performed the best and second best respectively (Figure 2A, Figure 2B). This ranking was consistent across sets of GO terms with different annotation frequencies, as well as for GO terms within each of the three GO ontology domains: Biological Process, Cellular Component, Molecular Function (Table S3). This ranking was also consistent when performance was calculated in different ways. Random Forest exhibited the best performance based on calculations of precision (proportion of predicted genes that are truly positive), recall (proportion of true positive genes recovered), F-measure (harmonic mean of precision and recall), consistency score, and AUC-ROC (Area Under Curve-Receiver Operator Curve) (Figure 2B). Ensemble methods were also evaluated however these did not show a significant increase in prediction accuracy compared to pure random forest based prediction (Figure S3).

**Figure 2:**
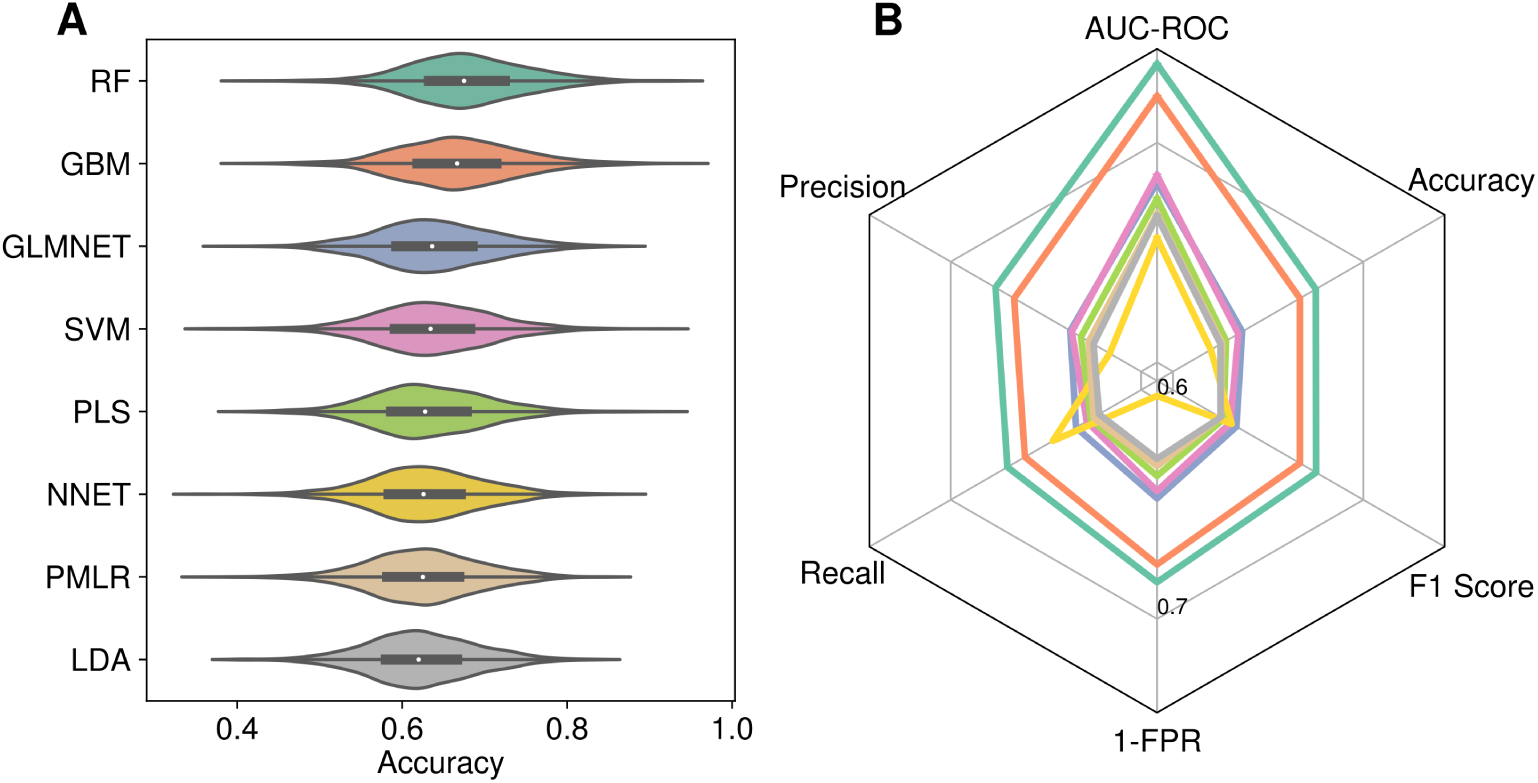
Performance of eight machine learning-based supervised classification algorithms in predicting gene functions using non-homology-based predictor variables. (A) Distribution of prediction accuracies for 1,562 GO terms using 8 methods. RF (Random Forest); GBM (Gradient Boosting Machine); GLMNET (Lasso and Elastic-Net Regularized Generalized Linear Models); SVM (Support Vector Machines with Radial Basis Function Kernel); PLS (Partial Least Squares); NNET (Neural Network); PMLR (Penalized Multinomial Logistic Regression); LDA (Linear Discriminant Analysis). (B) Median values for each of the eight algorithms. Color labeling in panel B correspond to the color labeling of each algorithm in panel A.

Among GO terms tested, the smallest class (GO terms assigned to 100-300 maize gene models) exhibits statistically significantly higher prediction accuracy (median= 0.688, SD=0.082) than the 300-600(median=0.688, SD=0.061), 600-900 (median=0.675, SD=0.053), 900-1200 (median=0.663, SD=0.047), 1200-1500 (median=0.661, SD=0.046), 1500-2000 (median=0.660, SD=0.041) or 2000-5000(median=0.641, SD=0.033) size classes (p=0.01, 2.54e-5, 4.91e-5, 5.57e-5, 6.29e-14, respectively; two tailed t-test) (Figure 3C, Table S3).

**Figure 3:**
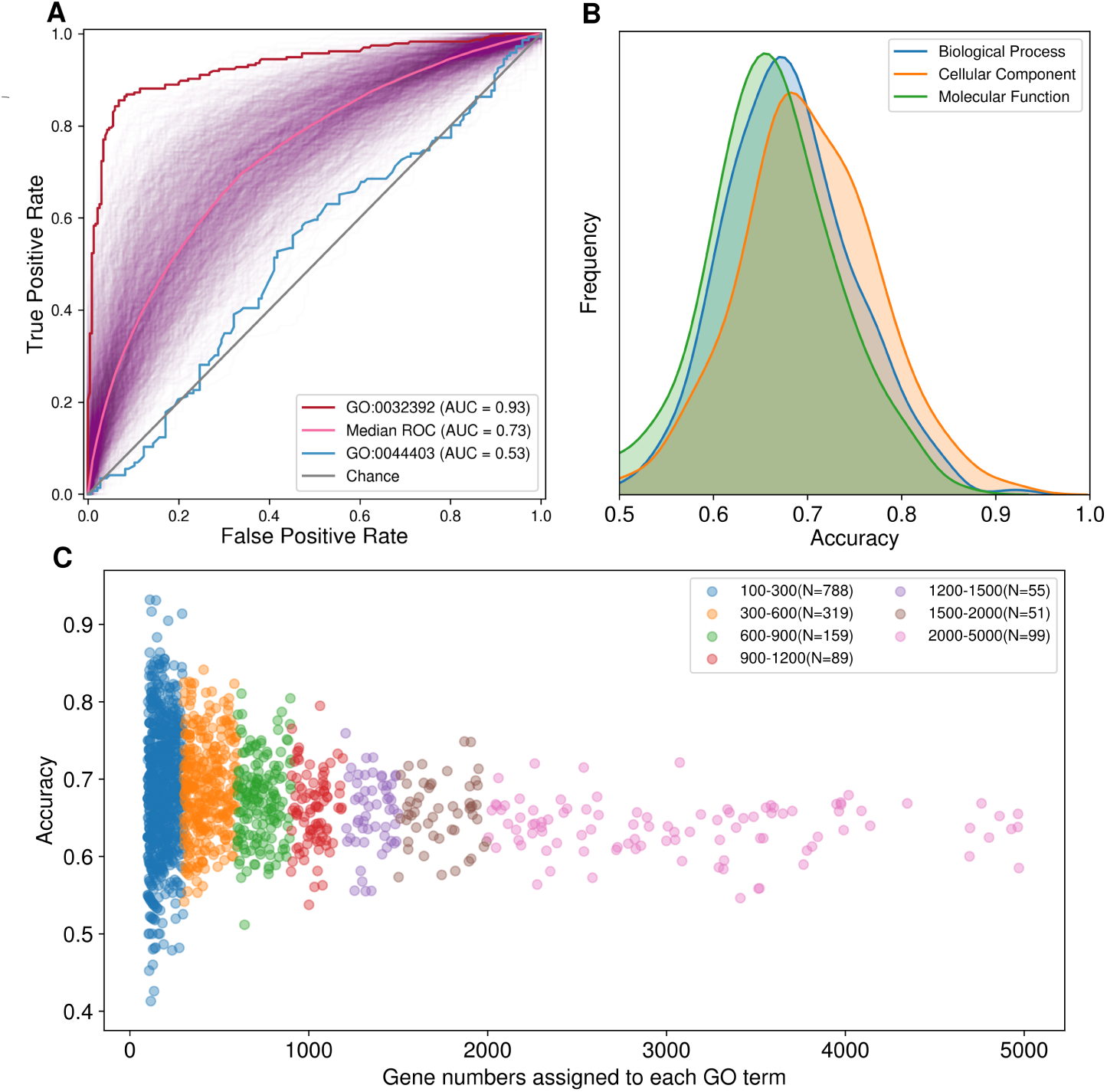
Prediction accuracy for individual GO terms varies in response to different characteristics of those terms. (A) Distribution of AUC-ROC values for random forest-based prediction of 1,562 GO terms, including information on the single best and second worst performing GO terms based on AUC-ROC (GO:0032392; DNA geometric change; Biological Process) and (GO:0044403; Symbiotic Process; Biological Process). The worst performing GO term (GO:005877) was for a biological process which does not occur in plants. (B) Distribution of prediction accuracies for individual GO terms in the Biological Process, Cellular Component and Molecular Function domains using random forest based prediction. (C) Relationship between prediction accuracy and the number of genes to which a given GO term is assigned in the Maize GAMER dataset.

### Higher Prediction Accuracy for Biological Process GO Terms

AUC-ROC values (area under curve-receiver operating characteristic) values were calculated for each individual GO term ROC (receiver operating characteristic) curves for the prediction accuracy of random forest based prediction – determined from 10-fold cross validation – were plotted for individual GO terms (Figure 3A). Details for the performance measures of every GO term provided in Table S4. As a control, AUC-ROC values were also calculated for genes with shuffled functional annotations. The 5^th^ and 95^th^ percentile of AUC-ROC values from 4 times of gene label shuffling for all GO terms were 0.45 and 0.56 with a median of 0.5. These values were consistent with expectations for random labeling of balanced data.

Random forest testing accuracy for individual GO terms ranged from 0.41 to 0.93 with a median of 0.68. The single best performing GO term prediction model, assessed based on accuracy, was for GO:0006270 (replication initiation) using random forest (precision = 95.2%, recall = 90.9%, FPR = 4.5%, Accuracy = 0.93, AUC-ROC = 0.92, Consistency score = 0.87). GO terms related to DNA replication (GO:0006270, DNA replication initiation, Accu = 0.93), modification (GO:0016556, mRNA modification, Accu = 0.90; GO:0006304, DNA modification, Accu = 0.85), methylation (GO:0006346, methylation-dependent chromatin silencing, Accu = 0.93; GO:0001510, RNA methylation, Accu = 0.91) and metabolic process (GO:0009220, pyrimidine ribonucleotide biosynthetic process, Accu = 0.86) are well predicted using nonhomology features. On the other end of the distribution, examples of GO terms with the lowest prediction accuracy were (GO:0022832, voltage-gated channel activity, Accu = 0.48; GO:0005216, ion channel activity, Accu = 0.48) and regulation of a process (GO:0050778, positive regulation of immune response, Accu = 0.52; GO:0051348, negative regulation of transferase activity). GO terms with higher prediction accuracy were drawn primarily from the Biological Process domain while GO terms with the lowest prediction accuracy belonged primarily to the Molecular Function domain.

To test whether this finding represented a consistent pattern, the distribution of prediction accuracies was evaluated separately for GO terms belonging to each of the three domains (Biological Process, Cellular Component, and Molecular Function). GO terms involved in Cellular Component have the highest median accuracy (Figure 3B). Cellular Component GO terms were the rarest of the three domains (151 GO terms of 1,562 total terms tested). Median accuracy for Biological Process GO terms was modestly lower than for Cellular Component. Biological Process GO terms were much more abundant (73%) in 1,562 GO terms test, which may explain why the most accurate individual GO terms were drawn from this domain. GO terms from the Molecular Function domain had the lowest median accuracy, and there were many Molecular Function GO terms, particularly those related to channel, transporter, enzyme activity or binding with extremely low accuracy (Table S4). This ranking of accuracy across GO domains was largely consistent across GO terms with different population sizes of genes carrying the annotation and across the results from predicting using different machine learning algorithms (Table S3).

### Contribution of Different Feature Types to Prediction Accuracy

Separate predictions were conducted using distinct subsets of features to assess relative contributions of different types of features to the overall accuracy of non-homology-based functional prediction. The ranking of prediction accuracy was largely consistent across the three primary GO term domains: Biological Process, Cellular Component, and Molecular Function. Models trained using only gene model structure features or trained using only RNA expression features provided approximately equal independent prediction accuracy. One exception was in the Molecular Function domain where models trained using only gene structure features performed almost equivalently to the complete model (AUC-ROC = 0.69 and 0.70 for the models using structural data only and full models, respectively). Models for predicting Molecular Function GO terms trained using only RNA expression features performed significantly worse than the complete model. Models trained using only chromatin features or only co-expression features did not perform well in any of the three domains (Figure 4A). However, at the level of individual GO terms, there were a number of GO terms where models trained using only protein expression features (99 GO terms 6.3%) or chromatin state features (35 GO terms 2.2%) had better performance than any of the other component models (Figure 4B). In a minor number of cases (15 GO terms 0.96%) the model trained using only co-expression features provided the highest accuracy of any of the component models (Table S5).

**Figure 4:**
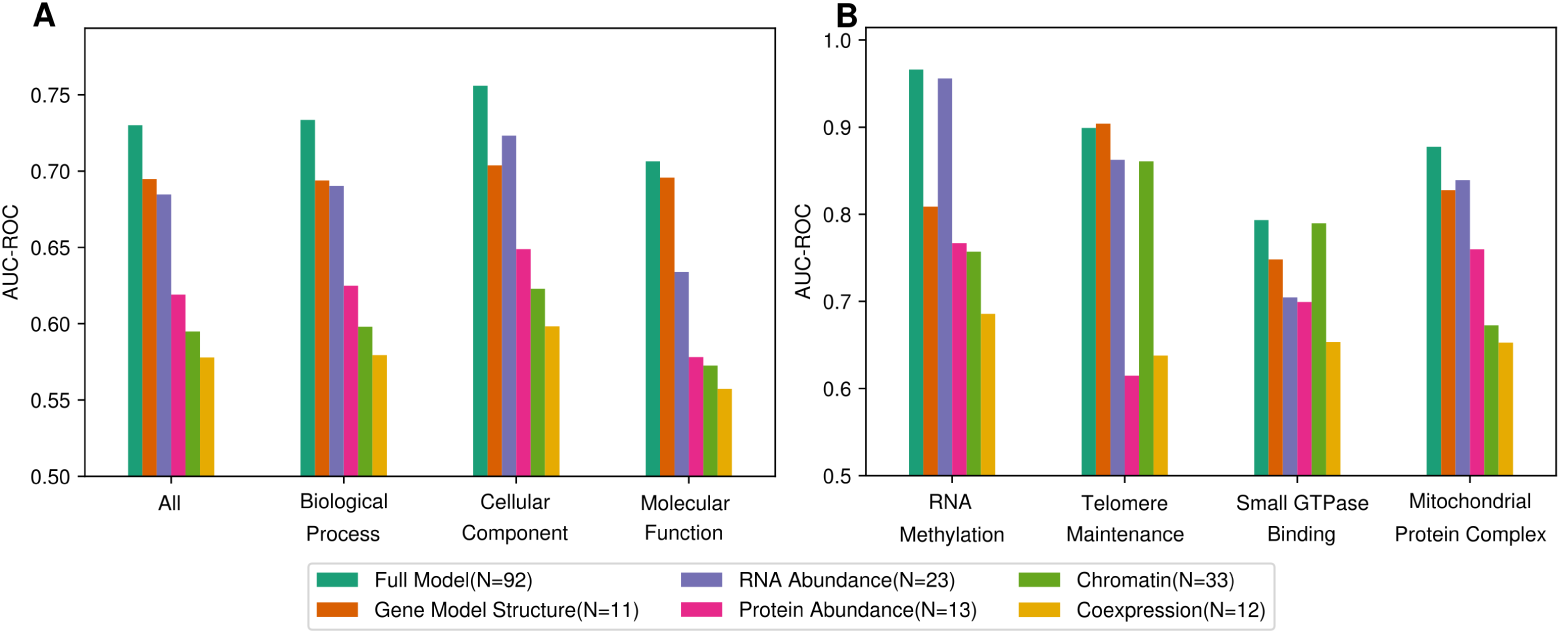
Contributions of each of five types of features to the overall functional prediction accuracy. (A) Median AUC-ROC values for all GO terms, and GO terms classified based on domain for the complete model, and partial models trained using only RNA abundance features, only chromatin features, only gene structure futures, only protein abundance features, or only co-expression features. (B) Examples of the same comparison shown in panel A for individual GO terms: RNA Methylation (GO:0001510) (Biological Process); Telomere Maintenance (GO:0000723) (Biological Process); Small GTPase Binding (GO:0031267) (Molecular Function); Mitochondrial Protein Complex (GO:0098798) (Cellular Component).

## Discussion

Accurate and precise annotation of gene model functions in the absence of gene by gene genetic analysis remains challenging. In most species, the vast majority of genes have not been studied or characterized directly. Instead, when functional annotations are present, they are drawn from functional characterization of homologous genes. Homology-based approaches may also introduce erroneous and misleading functional annotations. Firstly, genes which are homologous will not always perform the same biological function or be localized to the same cellular compartments. For example, R2R3-MYB transcription factors are all homologous to each other yet play different roles regulating responses to multiple stress conditions, controlling plant development and cell fate, or regulating secondary metabolism (Du et al., 2012). Secondly, because homology-based functional annotations are often drawn from datasets and databases which were originally also annotated based on homology, it is possible for incorrect functional annotations to propagate through biological databases indefinitely. Estimates of the mis-annotation using experimentally well-characterized sets of enzyme can range from about 25% to over 60% (Schnoes et al., 2009). Finally, 5% to 15% of annotated gene models in the genomes of many species are “orphans” without detectable homology to any protein with a characterized function. Here we sought to evaluate whether using machine learning methods and a set of non-homology-based features can complement existing methods for functional annotation. Non-homology-based methods may ultimately be able to correctly assign new functional annotations to gene models and identify potentially inaccurate existing functional annotations.

It is important to discuss one critical limitation of the analyses employed here. While non-homology-based annotation approaches ultimately hold the potential to identify and correct errors introduced by homology-based annotation, in this study a set of functional annotations derived from homology-based annotation were treated as ground truth. As the result, the true recall of non-homology-based methods may be higher than the estimated recall in this study, as some false negatives may in fact represent errors in the underlying functional annotations. At the same time, models trained using functional annotations currently assigned to only one or several homologous gene families may learn signatures of those gene families rather than the annotated function itself. In these cases, the accuracies calculated may be higher than the true values (Washburn et al., 2019). Going forward, there is a clear need for curated sets of experimentally supported functional annotations for maize equivalent to those previously generated for species such as yeast and arabidopsis (Aslett and Wood, 2006; Lamesch et al., 2011). While acknowledging these limitations some intriguing initial patterns are still apparent in this initial trial of non-homology based function annotation.

Machine learning-based functional annotation showed strengths which are complementary to known accuracy patterns of primarily homology-based methods. Specifically, homology-based functional annotation has been reported to show higher accuracy for GO terms in the Molecular Function domain (Radivojac et al., 2013; Jiang et al., 2016). In contrast, we found that non-homology-based predictions exhibited the highest prediction accuracy in the Cellular Component and Biological Process domains, and the lowest accuracy in Molecular Function (Figure 3B). Molecular functions (e.g. transcription factors, transporters, structural proteins) are likely to be conserved between homologous sequences. In contrast, the cellular localization and biological role of a given transcription factor or signal transduction component can vary and diverge substantially between even closely related homologs (Du et al., 2012). Genes involved in the same biological process or localized to a specific cellular compartment may be more likely to exhibit shared features such as co-expression than specific classes of transcription factors or transporters which may be localized to different cell types or expressed only in response to different environmental stimuli.

Going forward there are a number of potential avenues to improve the accuracy of genome-wide non-homology-based functional annotation. As discussed above, the incorporation of more detailed provenance information for existing functional annotations will serve both to train more accurate models, and to more accurately quantify the performance of these models. There are also additional types of non-homology-based predictive variables which could be incorporated in the future. These include more extensive protein and mRNA expression data, particularly from different stress conditions, experimentally derived protein-protein interaction data, descriptors of population genetic features including different types of selection and diversity, and as well as incorporating the results of quantitative genetic analyses using different types of phenotypes in different environments. Two challenges for future studies are how to integrate these heterogeneous data sources and how to deal with incomplete and noisy data.

## Methods

### Composition of the Prediction Variable Dataset

Predictive variables were divided into five categories: Gene Model Structure, RNA Expression, Protein Expression, Chromatin, Co-Expression, and Natural Diversity. Gene structural features included gene length from transcription start site to transcription stop site, including introns, exon number, CDS length, 3’ UTR and 5’ UTR length. These values were calculated for each gene using the published AGPv4 maize genome sequence and annotation (Jiao et al., 2017). Nucleotide composition and the GC content were calculated using all sequence from the annotated transcription start site to the annotated transcription stop site.

For protein-coding genes, a codon usage bias score which describes the degree of bias towards the most frequently used codons for multiple encoding amino acids in a given species was calculated following the method described in (Sharp and Li, 1987) as implemented in the SeqIO module in biopython (v1.72) package (Cock et al., 2009).

The initial set of RNA expression features included data from 2-3 replicates of 79 distinct tissue types in the maize inbred B73 (222 total samples) (Stelpflug et al., 2016) and 52 samples from biotic and biotic stress studies of B73 in different labs (Opitz et al., 2014; Makarevitch et al., 2015; Swart et al., 2017) for a total of 274 distinct samples. Normalized (FPKM: fragments per kilobase of exon per million aligned reads) expression values for each gene in each experiment were obtained from (Hoopes et al., 2019).

Protein expression features consisted of normalized protein abundance data quantified in dNSAF (distributed normalized spectral abundance factor) for 33 distinct tissues sampled from B73 were obtained from (Walley et al., 2016). B73 AGPv2 gene models were converted to B73 AGPv4 using a conversion list published on MaizeGDB (https://www.maizegdb.org/search/gene/downloadgenexrefs.php?relative=v4)

Chromatin features included DNA methylation (quantified separately in CG, CHG, and CHH contexts), three histone modifications (H3K4me3, H3K27me3, H3K27ac), and open chromatin as quantified by ATAC-seq. Raw sequence data for bisulfite-seq, ChIP-seq for H3K4me3, H3K27me3, and H3K27ac histone modifications, and ATAC-seq was downloaded from PRJNA391551 in the NCBI SRA (Dong et al., 2017). DNA methylation was quantified using Bismark (v0.19) with parameters “-L 50, -N 1” (Krueger and Andrews, 2011). ATAC-Seq and histone ChIP-seq reads were aligned to AGPv4 of the maize reference genome using gsnap (v2018-03-25) (Wu et al., 2016) with parameters “-m 0.02, -B 5, -n 1, -Q, –nofails”. Alignment files were then used to call peaks using the protocol previously described in (Dong et al., 2017).

For each of these chromatin features, scores were calculated for three regions: one using the gene body, defined as the region from the annotated transcription start site to the annotated transcription stop site, a second for the upstream region, defined as a 2 KB region directly upstream of the transcription start site, and a third for the downstream region, defined as the 2 KB region directly downstream of the transcription stop site. For each BS-seq dataset, for each of the three regions relative to each gene and each of three methylation contexts (CG, CHG, CHH), a single percentage score was calculated. This percentages was calculated as the ratio of all cytosines in that context in that genomic interval which were classified as “methylated” (≥ 5 mapped reads and with >50% of mapped reads showing methylation) to the total number of cytosines in that context in that genomic interval. For each ChIP-seq and ATAC-seq dataset, two features were calculated for each genomic interval: the maximum intensity among peaks overlapping that interval and the proportion of that interval covered by peaks using methods previously described in (Lloyd et al., 2017).

For each of these chromatin features, three scores were calculated: one using the gene body, defined as the region from the annotated transcription start site to the annotated transcription stop site, a second for the upstream region, defined as a 2KB region directly upstream of the transcription start site, and a third for the downstream region, defined as the 2 KB region directly downstream of the transcription stop site.

The co-expression set of features consisted of 12 binary variables defining membership in each of the 12 co-expression models defined by (Hoopes et al., 2019).

For population features, raw genotype calls of 277 resequenced inbreds in maize 282 association panel (Bukowski et al., 2017) were downloaded from Panzea (https://www.panzea.org/). Only biallelic SNPs were considered as variations in the given population for this study. SNP filtering, imputation and assignment to maize AGPv4 gene body region were processed as a previous study (Liang et al., 2019). SNP number per gene was determined by the number of final detected SNPs per AGPv4 gene.

### Dimension Reduction

Principal component-based dimension reduction was evaluated for RNA abundance and protein abundance data using R prcomp() function with parameters “center=TRUE, scale=TRUE”. For each set of features, 50 principal components were calculated. In each case, the decision on how many principal components to include was based on the cumulative proportion of variance explained.

### Defining the Subset of Gene Models and Functional Annotations

Several important features like protein abundance data for maize vegetative and reproductive stages, are only available for maize AGPv2. As the result, we constrained this analysis and only considered a set gene models which had a 1:1 relationship between a single gene model in the maize reference genome version AGPv2 and a single gene model in the maize reference genome version AGPv4 (Liang et al., 2019). A small number of genes with missing values for more than half of the total set of 369 features were omitted from subsequent analyses. For the remaining genes, features were centered, scaled and imputed (for missing values) using preProcess() function in caret (v6.0-80) R package (Kuhn, 2015).

An implicit GO term assignment can occur when a specific GO term is explicitly assigned to a gene, each parent of that GO term is also implicitly assigned to the same gene. The get all parents() function in goatools (v0.8.9) python package was used to add the implicit GO terms to each gene (Klopfenstein et al., 2018). After explicitly assigning implicit GO annotations to genes, GO terms which were assigned to less than 100 genes or more than 5,000 gene models were excluded.

### Implementing Machine Learning Algorithms

The eight machine learning algorithms, i.e. random forest, neural network, radial kernel SVM, glmnet, lda, penalized multinomial regression, partial least squares and gbm with parameter “tuneLength=5”, evaluated as part of this study were all implemented in the R package caret (v6.0-80) (Kuhn, 2015). For each GO term, a balanced training data was constructed using the set of maize genes assigned with that annotation as the “positive” set and a randomly selected equal number of genes not assigned with that annotation as the “negative” set. 20% of the negative and positive genes from each training set were set aside as the hold-out testing data to assess model performance. The remaining 80% of data was used to train each algorithm for each GO term. 10-fold cross validation was used. The three stacking ensemble methods evaluated in our study were also tested using implementations in the caret package. Each of the three was employed as a supervisor model and was provided with the output of three primary predictive methods (random forest, gbm, and glm) with “tuneLength=3”.

### Evaluating Prediction Accuracy

Accuracy, FPR (false positive rate), recall, precision, F1 score, AUC-ROC, and consistency score were calculated. Accuracy = (TP+TN)/(TP+TN+FP+FN) (where: TP = True positive; FP = False positive; TN = True negative; FN = False negative). The FPR was calculated as the ratio between the number of negative events wrongly categorized as positive and the total number of actual negative events (FP/FP+TN). Recall was defined as the fraction of relevant instances that have been retrieved over the total amount of relevant instances (TP/TP+FN). Precision was defined as the fraction of relevant instances among the retrieved instances (TP/TP+FP). F1 score was calculated as the harmonic mean of precision and recall. AUC-ROC was calculated as the ratio of the total the area under the plot of receiver operating characteristic curve, to the total area contained within the plot. For permutation testing to evaluate the potential of over-fitting, the same training and testing datasets were used and the same algorithms employed, but genes were shuffled between the positive and negative categories.

## Acknowledgements

This work was funded in part by the National Science Foundation under Grant Nos. (MCB-1838307 and OIA-1826781) to JCS, the Foundation for Food and Agricultural Research under grant No. 094525-17308 to JCS, National Science Foundation of China under grant No. 31871313 and Taishan Pandeng plan to PL. XD was supported by the a graduate fellowship from “Double-First Class” construction plan awarded by Shandong Agricultural University. This project was completed utilizing the Holland Computing Center of the University of Nebraska, which receives support from the Nebraska Research Initiative. The authors thank Daniel Carvalho for advice and guidance on the calculation of some potential predictor variables and Kevin Childs for providing access to maize salt stress expression data.

---

Table S1: Values for each gene for each of 369 features used in this study (TableS1-features-original-369.xlsx)

Table S2: List of all 369 and 92 features in the order they are show in Figure 1 (TableS2-feature-names.xlsx)

Table S3: Prediction performance for different algorithms on GO terms with different population sizes from different GO domains. (TableS3-population-size-domains.xlsx)

Table S4: Individual prediction performance each of the 1,562 GO terms used in this study.(TableS4-performances-summary.xlsx)

Table S5: AUC-ROC values for models trained with subsets of data (TableS5-AUC-summary-feature-imp.xlsx)

**Figure S1:**
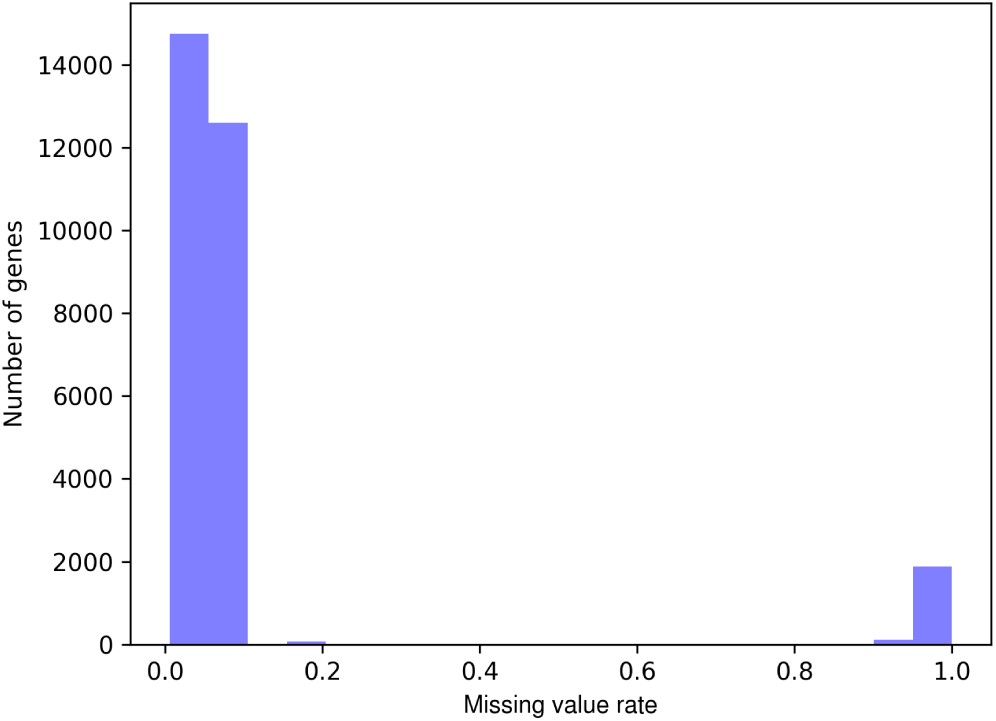
Distribution of missing rates, i.e. the proportion of 369 features for which a given gene has no reported value, among the initial set of 29,428 gene models with 1:1 matches between maize RefGen2 and maize RefGen4. Genes with high missing rates were excluded from downstream analyses.

**Figure S2:**
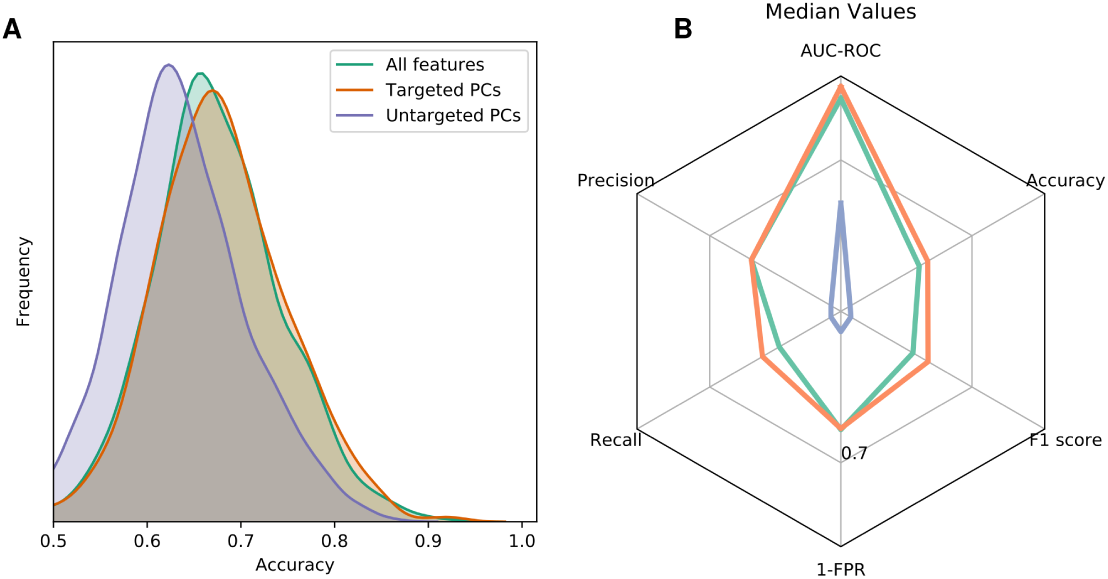
Comparison of prediction accuracy for models trained with sets of features after dimension reduction in different ways. (A) Distribution of prediction accuracies across all GO terms for random forest based predictions using either a set of all 369 features, a set of features with targeted and separate dimension reduction of mRNA and protein expression features, and a set of features generated by untargeted dimension-reduction of all 369 initial features into a set of principal components. (B) Median values for six different criteria scores of model performance for the same random forest based predictions shown in panel A. Color coding in panel B is the same as the legend shown in panel A.

**Figure S3:**
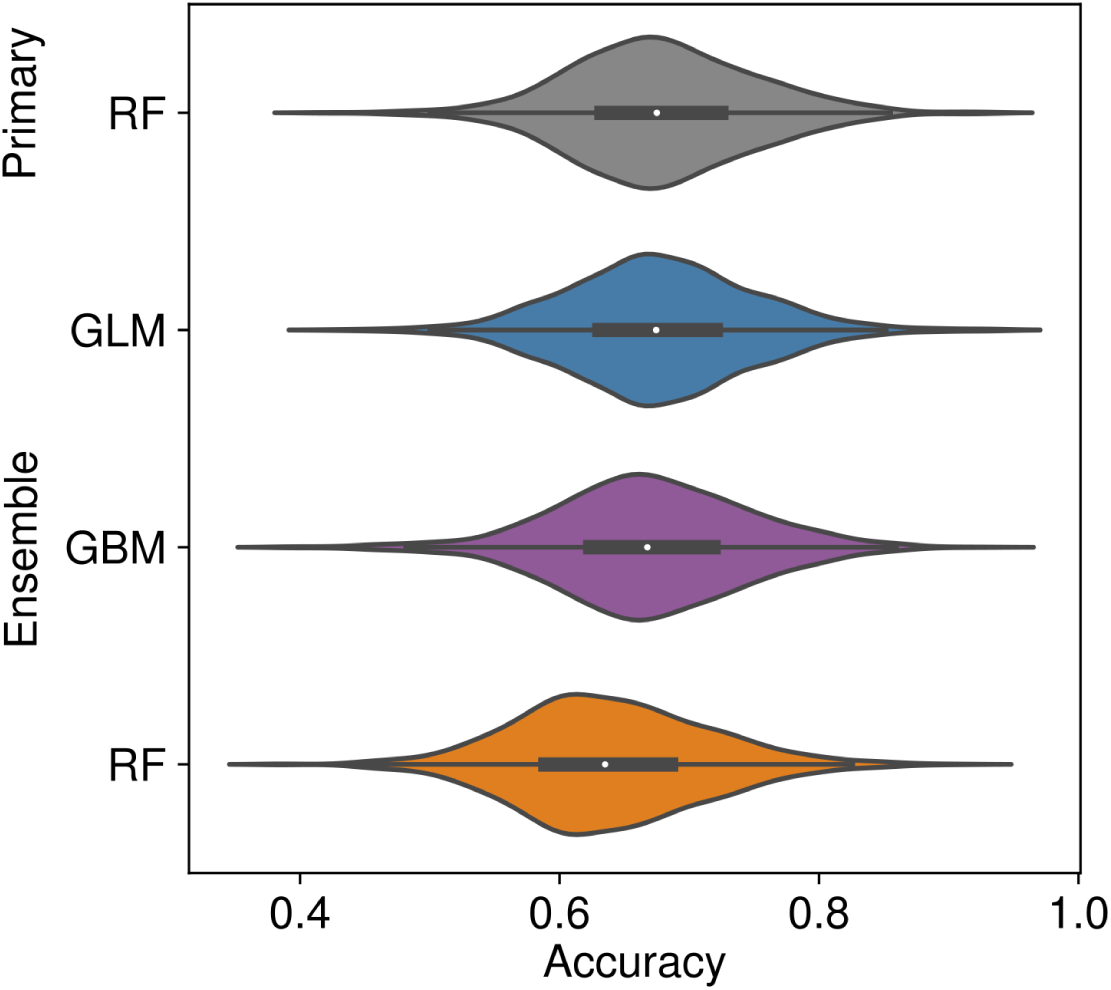
Distribution of prediction accuracies in all 1,562 tested GO terms for 3 ensemble methods aggregating the outputs from the three best-performing individual models, compared to the best-performing primary method (Random Forest) on its own. Abbreviations for the three ensemble methods are GBM (Gradient Boosting Machine); GLM (Generalized Linear Model); RF (Random Forest).

## References

Angelovici, R., Batushansky, A., Deason, N., Gonzalez-Jorge, S., Gore, M. A., Fait, A., and DellaPenna, D. (2017). Network-guided gwas improves identification of genes affecting free amino acids. Plant physiology, 173(1):872–886.

Aslett, M. and Wood, V. (2006). Gene ontology annotation status of the fission yeast genome: preliminary coverage approaches 100%. Yeast, 23(13):913–919.

Baldauf, J. A., Marcon, C., Lithio, A., Vedder, L., Altrogge, L., Piepho, H.-P., Schoof, H., Nettleton, D., and Hochholdinger, F. (2018). Single-parent expression is a general mechanism driving extensive complementation of non-syntenic genes in maize hybrids. Current Biology, 28(3):431–437.

Baldauf, J. A., Marcon, C., Paschold, A., and Hochholdinger, F. (2016). Nonsyntenic genes drive tissue-specific dynamics of differential, nonadditive, and allelic expression patterns in maize hybrids. Plant physiology, 171(2):1144–1155.

Brenner, S. E. (1999). Errors in genome annotation. Trends in Genetics, 15(4):132–133.

Brown, S. D., Gerlt, J. A., Seffernick, J. L., and Babbitt, P. C. (2006). A gold standard set of mechanistically diverse enzyme superfamilies. Genome biology, 7(1):R8.

Bukowski, R., Guo, X., Lu, Y., Zou, C., He, B., Rong, Z., Wang, B., Xu, D., Yang, B., Xie, C., et al. (2017). Construction of the third-generation zea mays haplotype map. Gigascience, 7(4):gix134.

Campbell, M. S., Law, M., Holt, C., Stein, J. C., Moghe, G. D., Hufnagel, D. E., Lei, J., Achawanantakun, R., Jiao, D., Lawrence, C. J., et al. (2014). Maker-p: a tool kit for the rapid creation, management, and quality control of plant genome annotations. Plant physiology, 164(2):513–524.

Chan, E. K., Rowe, H. C., Corwin, J. A., Joseph, B., and Kliebenstein, D. J. (2011). Combining genome-wide association mapping and transcriptional networks to identify novel genes controlling glucosinolates in arabidopsis thaliana. PLoS biology, 9(8):e1001125.

Chen, F., Dong, W., Zhang, J., Guo, X., Chen, J., Wang, Z., Lin, Z., Tang, H., and Zhang, L. (2018). The sequenced angiosperm genomes and genome databases. Frontiers in plant science, 9:418.

Clark, W. T. and Radivojac, P. (2011). Analysis of protein function and its prediction from amino acid sequence. Proteins: Structure, Function, and Bioinformatics, 79(7):2086–2096.

Cock, P. J., Antao, T., Chang, J. T., Chapman, B. A., Cox, C. J., Dalke, A., Friedberg, I., Hamelryck, T., Kauff, F., Wilczynski, B., et al. (2009). Biopython: freely available python tools for computational molecular biology and bioinformatics. Bioinformatics, 25(11):1422–1423.

Conesa, A., Götz, S., García-Gómez, J. M., Terol, J., Talón, M., and Robles, M. (2005). Blast2go: a universal tool for annotation, visualization and analysis in functional genomics research. Bioinformatics, 21(18):3674–3676.

Cook, D. E., Valle-Inclan, J. E., Pajoro, A., Rovenich, H., Thomma, B. P., and Faino, L. (2019). Long-read annotation: automated eukaryotic genome annotation based on long-read cdna sequencing. Plant physiology, 179(1):38–54.

Del Angel, V. D., Hjerde, E., Sterck, L., Capella-Gutierrez, S., Notredame, C., Pettersson, O. V., Amselem, J., Bouri, L., Bocs, S., Klopp, C., et al. (2018). Ten steps to get started in genome assembly and annotation. F1000Research, 7.

Dong, P., Tu, X., Chu, P.-Y., Lü, P., Zhu, N., Grierson, D., Du, B., Li, P., and Zhong, S. (2017). 3d chromatin architecture of large plant genomes determined by local a/b compartments. Molecular plant, 10(12):1497–1509.

Du, H., Feng, B.-R., Yang, S.-S., Huang, Y.-B., and Tang, Y.-X. (2012). The r2r3-myb transcription factor gene family in maize. PloS one, 7(6):e37463.

Edwards, M. T., Rison, S. C., Stoker, N. G., and Wernisch, L. (2005). A universally applicable method of operon map prediction on minimally annotated genomes using conserved genomic context. Nucleic acids research, 33(10):3253–3262.

Enault, F., Suhre, K., and Claverie, J.-M. (2005). Phydbac” gene function predictor”: a gene annotation tool based on genomic context analysis. BMC bioinformatics, 6(1):247.

Finn, R. D., Coggill, P., Eberhardt, R. Y., Eddy, S. R., Mistry, J., Mitchell, A. L., Potter, S. C., Punta, M., Qureshi, M., Sangrador-Vegas, A., et al. (2015). The pfam protein families database: towards a more sustainable future. Nucleic acids research, 44(D1):D279–D285.

Friedman, J., Hastie, T., and Tibshirani, R. (2010). Regularization paths for generalized linear models via coordinate descent. Journal of Statistical Software, 33(1):1–22.

Gaasterland, T. and Ragan, M. A. (1998). Microbial genescapes: phyletic and functional patterns of orf distribution among prokaryotes. Microbial & comparative genomics, 3(4):199–217.

Gabaldón, T. and Huynen, M. A. (2004). Prediction of protein function and pathways in the genome era. Cellular and Molecular Life Sciences CMLS, 61(7-8):930–944.

Gilks, W. R., Audit, B., De Angelis, D., Tsoka, S., and Ouzounis, C. A. (2002). Modeling the percolation of annotation errors in a database of protein sequences. Bioinformatics, 18(12):1641–1649.

Gilks, W. R., Audit, B., de Angelis, D., Tsoka, S., and Ouzounis, C. A. (2005). Percolation of annotation errors through hierarchically structured protein sequence databases. Mathematical biosciences, 193(2):223–234.

Goodstein, D. M., Shu, S., Howson, R., Neupane, R., Hayes, R. D., Fazo, J., Mitros, T., Dirks, W., Hellsten, U., Putnam, N., et al. (2011). Phytozome: a comparative platform for green plant genomics. Nucleic acids research, 40(D1):D1178–D1186.

Guo, W.-J., Li, P., Ling, J., and Ye, S.-P. (2007). Significant comparative characteristics between orphan and nonorphan genes in the rice (oryza sativa l.) genome. International Journal of Genomics, 2007.

Guo, Y.-L. (2013). Gene family evolution in green plants with emphasis on the origination and evolution of a rabidopsis thaliana genes. The Plant Journal, 73(6):941–951.

Hoopes, G. M., Hamilton, J. P., Wood, J. C., Esteban, E., Pasha, A., Vaillancourt, B., Provart, N. J., and Buell, C. R. (2019). An updated gene atlas for maize reveals organ-specific and stress-induced genes. The Plant Journal, 97(6):1154–1167.

Hulo, N., Bairoch, A., Bulliard, V., Cerutti, L., De Castro, E., Langendijk-Genevaux, P. S., Pagni, M., and Sigrist, C. J. (2006). The prosite database. Nucleic acids research, 34(suppl 1):D227–D230.

Iyer, L. M., Aravind, L., Bork, P., Hofmann, K., Mushegian, A. R., Zhulin, I. B., and Koonin, E. V. (2001). Quod erat demonstrandum? the mystery of experimental validation of apparently erroneous computational analyses of protein sequences. Genome Biology, 2(12):research0051–1.

Jiang, Y., Oron, T. R., Clark, W. T., Bankapur, A. R., D’Andrea, D., Lepore, R., Funk, C. S., Kahanda, I., Verspoor, K. M., Ben-Hur, A., et al. (2016). An expanded evaluation of protein function prediction methods shows an improvement in accuracy. Genome biology, 17(1):184.

Jiao, Y., Peluso, P., Shi, J., Liang, T., Stitzer, M. C., Wang, B., Campbell, M. S., Stein, J. C., Wei, X., Chin, C.-S., et al. (2017). Improved maize reference genome with single-molecule technologies. Nature, 546(7659):524.

Jones, C. E., Brown, A. L., and Baumann, U. (2007). Estimating the annotation error rate of curated go database sequence annotations. BMC bioinformatics, 8(1):170.

Karatzoglou, A., Smola, A., Hornik, K., and Karatzoglou, M. A. (2018). Package ‘kernlab’. Technical report, Technical report, CRAN, 03 2016.

Karatzoglou, A., Smola, A., Hornik, K., and Zeileis, A. (2004). kernlab – an S4 package for kernel methods in R. Journal of Statistical Software, 11(9):1–20.

Karp, P. D. (1998). What we do not know about sequence analysis and sequence databases. Bioinformatics (Oxford, England), 14(9):753–754.

Klopfenstein, D., Zhang, L., Pedersen, B. S., Ramírez, F., Vesztrocy, A. W., Naldi, A., Mungall, C. J., Yunes, J. M., Botvinnik, O., Weigel, M., et al. (2018). Goatools: A python library for gene ontology analyses. Scientific reports, 8(1):10872.

Krueger, F. and Andrews, S. R. (2011). Bismark: a flexible aligner and methylation caller for bisulfite-seq applications. bioinformatics, 27(11):1571–1572.

Kuhn, M. (2015). Caret: classification and regression training. Astrophysics Source Code Library.

Lamesch, P., Berardini, T. Z., Li, D., Swarbreck, D., Wilks, C., Sasidharan, R., Muller, R., Dreher, K., Alexander, D. L., Garcia-Hernandez, M., et al. (2011). The arabidopsis information resource (tair): improved gene annotation and new tools. Nucleic acids research, 40(D1):D1202–D1210.

Liang, Z., Qiu, Y., and Schnable, J. (2019). Distinct characteristics of genes associated with phenome-wide variation in maize (zea mays). bioRxiv, page 534503.

Liaw, A. and Wiener, M. (2002). Classification and regression by randomforest. R News, 2(3):18–22.

Lloyd, J. P., Tsai, Z. T., Sowers, R. P., Panchy, N. L., and Shiu, S.-H. (2017). Defining the functional significance of intergenic transcribed regions based on heterogeneous features of phenotype genes and pseudogenes. bioRxiv, page 127282.

Lock, A., Rutherford, K., Harris, M. A., Hayles, J., Oliver, S. G., Bähler, J., and Wood, V. (2018). Pombase 2018: user-driven reimplementation of the fission yeast database provides rapid and intuitive access to diverse, interconnected information. Nucleic acids research, 47(D1):D821–D827.

Makarevitch, I., Waters, A. J., West, P. T., Stitzer, M., Hirsch, C. N., Ross-Ibarra, J., and Springer, N. M. (2015). Transposable elements contribute to activation of maize genes in response to abiotic stress. PLoS genetics, 11(1):e1004915.

Marcotte, E. M. (2000). Computational genetics: finding protein function by nonhomology methods. Current opinion in structural biology, 10(3):359–365.

Marcotte, E. M., Pellegrini, M., Ng, H.-L., Rice, D. W., Yeates, T. O., and Eisenberg, D. (1999). Detecting protein function and protein-protein interactions from genome sequences. Science, 285(5428):751–753.

Michael, T. P. and Jackson, S. (2013). The first 50 plant genomes. The plant genome, 6(2).

Monnahan, P. J., Michno, J.-M., O’Connor, C. H., Brohammer, A. B., Springer, N. M., McGaugh, S. E., and Hirsch, C. N. (2019). Using multiple reference genomes to identify and resolve annotation inconsistencies. bioRxiv, page 651984.

Morett, E., Korbel, J. O., Rajan, E., Saab-Rincon, G., Olvera, L., Olvera, M., Schmidt, S., Snel, B., and Bork, P. (2003). Systematic discovery of analogous enzymes in thiamin biosynthesis. Nature biotechnology, 21(7):790.

Oellrich, A., Walls, R. L., Cannon, E. K., Cannon, S. B., Cooper, L., Gardiner, J., Gkoutos, G. V., Harper, L., He, M., Hoehndorf, R., et al. (2015). An ontology approach to comparative phenomics in plants. Plant methods, 11(1):10.

Opitz, N., Paschold, A., Marcon, C., Malik, W. A., Lanz, C., Piepho, H.-P., and Hochholdinger, F. (2014). Transcriptomic complexity in young maize primary roots in response to low water potentials. BMC genomics, 15(1):741.

Paschold, A., Larson, N. B., Marcon, C., Schnable, J. C., Yeh, C.-T., Lanz, C., Nettleton, D., Piepho, H.-P., Schnable, P. S., and Hochholdinger, F. (2014). Nonsyntenic genes drive highly dynamic complementation of gene expression in maize hybrids. The Plant Cell, 26(10):3939–3948.

Pellegrini, M., Marcotte, E. M., Thompson, M. J., Eisenberg, D., and Yeates, T. O. (1999). Assigning protein functions by comparative genome analysis: protein phylogenetic profiles. Proceedings of the National Academy of Sciences, 96(8):4285–4288.

Quevillon, E., Silventoinen, V., Pillai, S., Harte, N., Mulder, N., Apweiler, R., and Lopez, R. (2005). Interproscan: protein domains identifier. Nucleic acids research, 33(suppl 2):W116–W120.

Radivojac, P., Clark, W. T., Oron, T. R., Schnoes, A. M., Wittkop, T., Sokolov, A., Graim, K., Funk, C., Verspoor, K., Ben-Hur, A., et al. (2013). A large-scale evaluation of computational protein function prediction. Nature methods, 10(3):221.

Ridgeway, G., Southworth, M. H., and RUnit, S. (2013). Package ‘gbm’. Viitattu, 10(2013):40.

Ripley, B., Venables, B., Bates, D. M., Hornik, K., Gebhardt, A., Firth, D., and Ripley, M. B. (2013). Package ‘mass’. Cran R.

Ripley, B., Venables, W., and Ripley, M. B. (2016). Package ‘nnet’. R package version, 7:3–12.

Schaefer, R. J., Michno, J.-M., Jeffers, J., Hoekenga, O., Dilkes, B., Baxter, I., and Myers, C. L. (2018). Integrating coexpression networks with gwas to prioritize causal genes in maize. The Plant Cell, 30(12):2922–2942.

Schnable, J. C. and Freeling, M. (2011). Genes identified by visible mutant phenotypes show increased bias toward one of two subgenomes of maize. PloS one, 6(3):e17855.

Schnable, P. S., Ware, D., Fulton, R. S., Stein, J. C., Wei, F., Pasternak, S., Liang, C., Zhang, J., Fulton, L., Graves, T. A., et al. (2009). The b73 maize genome: complexity, diversity, and dynamics. science, 326(5956):1112–1115.

Schnoes, A. M., Brown, S. D., Dodevski, I., and Babbitt, P. C. (2009). Annotation error in public databases: misannotation of molecular function in enzyme superfamilies. PLoS computational biology, 5(12):e1000605.

Sharp, P. M. and Li, W.-H. (1987). The codon adaptation index-a measure of directional synonymous codon usage bias, and its potential applications. Nucleic acids research, 15(3):1281–1295.

Simon, N., Friedman, J., Hastie, T., and Tibshirani, R. (2011). Regularization paths for cox’s proportional hazards model via coordinate descent. Journal of Statistical Software, 39(5):1–13.

Springer, N. M., Anderson, S. N., Andorf, C. M., Ahern, K. R., Bai, F., Barad, O., Barbazuk, W. B., Bass, H. W., Baruch, K., Ben-Zvi, G., et al. (2018). The maize w22 genome provides a foundation for functional genomics and transposon biology. Nature genetics, 50(9):1282.

Springer, N. M., Ying, K., Fu, Y., Ji, T., Yeh, C.-T., Jia, Y., Wu, W., Richmond, T., Kitzman, J., Rosenbaum, H., et al. (2009). Maize inbreds exhibit high levels of copy number variation (cnv) and presence/absence variation (pav) in genome content. PLoS genetics, 5(11):e1000734.

Stelpflug, S. C., Sekhon, R. S., Vaillancourt, B., Hirsch, C. N., Buell, C. R., de Leon, N., and Kaeppler, S. M. (2016). An expanded maize gene expression atlas based on rna sequencing and its use to explore root development. The plant genome, 9(1).

Stoeger, T., Gerlach, M., Morimoto, R. I., and Amaral, L. A. N. (2018). Large-scale investigation of the reasons why potentially important genes are ignored. PLoS biology, 16(9):e2006643.

Sun, S., Zhou, Y., Chen, J., Shi, J., Zhao, H., Zhao, H., Song, W., Zhang, M., Cui, Y., Dong, X., et al. (2018). Extensive intraspecific gene order and gene structural variations between mo17 and other maize genomes. Nature genetics, 50(9):1289.

Swanson-Wagner, R. A., Eichten, S. R., Kumari, S., Tiffin, P., Stein, J. C., Ware, D., and Springer, N. M. (2010). Pervasive gene content variation and copy number variation in maize and its undomesticated progenitor. Genome research, 20(12):1689–1699.

Swart, V., Crampton, B. G., Ridenour, J. B., Bluhm, B. H., Olivier, N. A., Meyer, J. M., and Berger, D. K. (2017). Complementation of ctb7 in the maize pathogen cercospora zeina overcomes the lack of in vitro cercosporin production. Molecular plant-microbe interactions, 30(9):710–724.

Swigoňová, Z., Lai, J., Ma, J., Ramakrishna, W., Llaca, V., Bennetzen, J. L., and Messing, J. (2004). Close split of sorghum and maize genome progenitors. Genome research, 14(10a):1916–1923.

Tang, J., Alelyani, S., and Liu, H. (2014). Feature selection for classification: A review. Data classification: algorithms and applications, page 37.

Tello-Ruiz, M. K., Stein, J., Wei, S., Preece, J., Olson, A., Naithani, S., Amarasinghe, V., Dharmawardhana, P., Jiao, Y., Mulvaney, J., et al. (2015). Gramene 2016: comparative plant genomics and pathway resources. Nucleic acids research, 44(D1):D1133–D1140.

Thomas, P. D., Campbell, M. J., Kejariwal, A., Mi, H., Karlak, B., Daverman, R., Diemer, K., Muruganujan, A., and Narechania, A. (2003). Panther: a library of protein families and subfamilies indexed by function. Genome research, 13(9):2129–2141.

Valencia, A. (2005). Automatic annotation of protein function. Current opinion in structural biology, 15(3):267–274.

Venables, W. N. and Ripley, B. D. (2002). Modern Applied Statistics with S. Springer, New York, fourth edition. ISBN 0-387-95457-0.

Walley, J. W., Sartor, R. C., Shen, Z., Schmitz, R. J., Wu, K. J., Urich, M. A., Nery, J. R., Smith, L. G., Schnable, J. C., Ecker, J. R., et al. (2016). Integration of omic networks in a developmental atlas of maize. Science, 353(6301):814–818.

Wang, B., Tseng, E., Regulski, M., Clark, T. A., Hon, T., Jiao, Y., Lu, Z., Olson, A., Stein, J. C., and Ware, D. (2016). Unveiling the complexity of the maize transcriptome by single-molecule long-read sequencing. Nature communications, 7:11708.

Washburn, J. D., Mejia-Guerra, M. K., Ramstein, G., Kremling, K. A., Valluru, R., Buckler, E. S., and Wang, H. (2019). Evolutionarily informed deep learning methods for predicting relative transcript abundance from dna sequence. Proceedings of the National Academy of Sciences, 116(12):5542–5549.

Wehrens, R. and Mevik, B.-H. (2007). The pls package: principal component and partial least squares regression in r. Journal of Statistical Software, 18.

Wimalanathan, K., Friedberg, I., Andorf, C. M., and Lawrence-Dill, C. J. (2018). Maize go annotation—methods, evaluation, and review (maize-gamer). Plant Direct, 2(4):e00052.

Wu, T. D., Reeder, J., Lawrence, M., Becker, G., and Brauer, M. J. (2016). Gmap and gsnap for genomic sequence alignment: enhancements to speed, accuracy, and functionality. In Statistical Genomics, pages 283–334. Springer.

Zheng, J., He, C., Qin, Y., Lin, G., Park, W. D., Sun, M., Li, J., Lu, X., Zhang, C., Yeh, C.-T., et al. (2019). Co-expression analysis aids in the identification of genes in the cuticular wax pathway in maize. The Plant Journal, 97(3):530–542.

